# Awake Hippocampal-Cortical Co-reactivation Is Associated with Forgetting

**DOI:** 10.1101/2022.12.10.519896

**Authors:** Büşra Tanrıverdi, Emily T Cowan, Athanasia Metoki, Katie R. Jobson, Vishnu P. Murty, Jason Chein, Ingrid R. Olson

**Affiliations:** Department of Psychology and Neuroscience, Temple University, Philadelphia, PA, USA; Department of Neurology, Washington University, St. Louis MO, USA

**Author notes:** Address correspondence to: Büşra Tanrıverdi, Department of Psychology and Neuroscience, Temple University, 1701 N. 13^th^ Street, Philadelphia, PA 19122.

**Keywords:** Consolidation, co-reactivation, co-replay, episodic memory, forgetting, hippocampus, replay, reactivation

## Abstract

Systems consolidation theories posit that consolidation occurs primarily through a coordinated communication between hippocampus and neocortex (McClelland and O’Reilly 1995; Kumaran et al., 2016; Moscovitch and Gilboa, 2022). Recent sleep studies in rodents have shown that hippocampus and visual cortex replay the same information at temporal proximity (“co-replay”) (Ji & Wilson, 2007; Lansink et al., 2009; Wierzynski et al., 2009; Peyrache et al., 2009). We developed a novel TR-based co-reactivation (TRCR) analysis method to study hippocampal-cortical co-replays in humans using functional MRI. Thirty-six young adults completed an image (face or scene)-location paired associate encoding task in the scanner, which were preceded and followed by resting state scans. We identified post-encoding rest TRs (+/− 1) that showed neural reactivation of each image-location trials in both hippocampus (HPC) and category-selective cortex (fusiform face area, FFA). This allowed us to characterize temporally proximal coordinated reactivations (“co-reactivations”) between HPC and FFA. Moreover, we found that increased HPC-FFA co-reactivations were associated with incorrectly recognized trials after a 1-week delay (*p* = 0.004). Finally, we found that these HPC-FFA co-reactivations were also associated with trials that were initially correctly recognized immediately after encoding but were later forgotten in 1-day (*p* = 0.043) and 1-week delay period (*p* = 0.031). We discuss these results from a trace transformation perspective (Winocur and Moscovitch, 2011; Sekeres et al., 2018) and speculate that HPC-FFA co-reactivations may be integrating related events, at the expense of disrupting event-specific details, hence leading to forgetting.

## Introduction

For several decades, research in rodents has demonstrated that neuronal firing patterns present at learning are replayed by hippocampal cells during post-encoding sleep (Buzsaki, 1989; Wilson and McNaughton, 1994; Skaggs and McNaughton, 1996; Girardeau and Zugaro, 2011) and awake rest (Foster and Wilson, 2006; Diba and Buzsáki 2007; Davidson et al. 2009; Karlsson and Frank 2009; Jadhav et al. 2012), that preserve the spatiotemporal properties of previously learned representations. It is thought that neural reactivation, such as described in these studies, is a key memory consolidation mechanism that first acts to strengthen the encoding patterns within hippocampus (Rasch and Born 2007; Carr et al. 2011), and then gradually integrates new events with related representations stored in the cortex (Tambini & Davachi, 2019).

Systems Consolidation views of memory, such as the Complementary Learning Systems (CLS) model, emphasize that consolidation occurs primarily through a coordinated communication between hippocampus and neocortex (McClelland and O’Reilly 1995; Nadel and Moscovitch, 1997; Winocur and Moscovitch, 2011; Kumaran et al., 2016; Robin and Moscovitch, 2017), including sensory cortex. A small number of sleep studies in rodents have provided evidence for “coordinated replay” such that the hippocampus and category selective cortices replay the same information around the same time (Ji & Wilson, 2007; Lansink et al., 2009; Wierzynski et al., 2009; Peyrache et al., 2009).

Notably, in these types of models, the nature of representations across the hippocampus and cortex varies such that the hippocampus stores more veridical accounts of an event, whereas the cortex stores more schematic representations reflecting commonalities amongst similar events. These models leave open questions about how coordinated activation across the hippocampus and cortex relate to memory for unique events. Coordinated replay could strengthen detailed representations of the prior events or alternatively, it could bias representations towards commonalities and/or introduce noise into the reactivated representation thereby degrading the precise details of encoded events.

Similar to what has been observed in rodents, an emerging body of work using human functional MRI (fMRI) has reported post-encoding reactivation during awake rest (see Tambini and Davachi, 2019 for a review) and shown that such reactivation appears to be related to better memory. More recently, researchers have looked at event-specific reactivation in the human brain, using multivariate approaches to examine the similarity between patterns of brain activity during encoding and post-encoding rest periods (Staresina et al., 2013; Deuker et al., 2013; Schlichting & Preston, 2014; Alm et al., 2019). For instance, Staresina et al (2013) used representational similarity analysis (RSA) and found that human entorhinal cortex (ErC) showed greater reactivation of subsequently remembered object-scene pairs during post-encoding rest. However, others have found that weakly encoded (see Schapiro et al., 2018) or weakly attended events (Jafarpour et al., 2017) are prioritized for reactivation. Therefore, how awake reactivations in the human brain prioritize and consolidate information is still unclear in human brain, although common assumptions are made that these reactivations are beneficial for memory.

Notably, none of this prior work investigating event-specific reinstatement probed the role of sensory representations in conjunction with the hippocampus, precluding the ability to evaluate the consequence of more systems-like consolidation. The question of whether or how reactivations are coordinated between the hippocampus and cortex has not been directly addressed in human fMRI studies. Studies that investigated hippocampal-cortical interactions during awake post-encoding rest utilized connectivity as a proxy for ‘coordinated replay’ and reported greater functional connectivity between the hippocampus and areas of the cortex during post-compared to pre-encoding rest for subsequently remembered information (Tambini, Kertz and Davachi, 2010; Tompary, Duncan et al., 2015, Tompary et al., 2017; Gruber et al., 2016; Murty et al., 2017). To our knowledge, no human fMRI study has yet tested event-specific hippocampal-cortical coordinated reactivations (‘*co-reactivations*”).

Here we tested two hypotheses, derived from theories of systems consolidation: 1) the hippocampus and cortex should co-reactivate information and 2) this co-reactivation should correlate with subsequent memory performance. To assess these hypotheses in a more targeted way, we developed a novel TR-based co-reactivation (TRCR) method, providing a “proof of concept” that concurrent reactivation patterns in hippocampal and cortical sites can be identified using fMRI. To test this, while in the scanner, participants first completed a 6-min rest scan (*pre-encoding),* followed by two encoding blocks, which were each followed by a post-task rest scan (*post-encoding)* (Fig. 1A). During encoding, participants were shown images of one of two possible categories (face or scene, in separate blocks) paired with a unique location on a 4-by-4 grid (*image trials,* henceforth). Leveraging our design, our analyses focused on examining co-reactivation during the post-encoding rest periods between the hippocampus (HPC) and two cortical regions of interest (ROIs) in the fusiform face area (FFA) and parahippocampal place area (PPA). Cortical ROIs were a priori selected given their previously established role in processing category-selective information (e.g., faces in FFA, Kanwisher and Yovel, 2006; and scenes in PPA, Epstein and Kanwisher, 1998; Park and Chun, 2009). We defined co-reactivation as reactivation in HPC and cortex to the same image trial, at the same TR (+/− 1 TR; see Fig. 1B, C). We then examined the relationship between co-reactivations and subsequent memory based on recognition tasks conducted immediately upon leaving the scanner, 1-day, and 1-week after learning. On these memory tasks, participants were cued with a location on a 4-by-4 grid and were instructed to select the correct image paired with this location from among three image options (a target and two lures). Our analyses revealed that there were more hippocampal-cortical co-reactivations during post-than pre-encoding rest, establishing this novel TR-based co-reactivation (TRCR) approach as a successful method to study hippocampal-cortical interactions during the consolidation window. Moreover, we found that these hippocampal-cortical co-reactivations were uniquely associated with forgetting, posing questions about their functional role.

**Fig 1.**
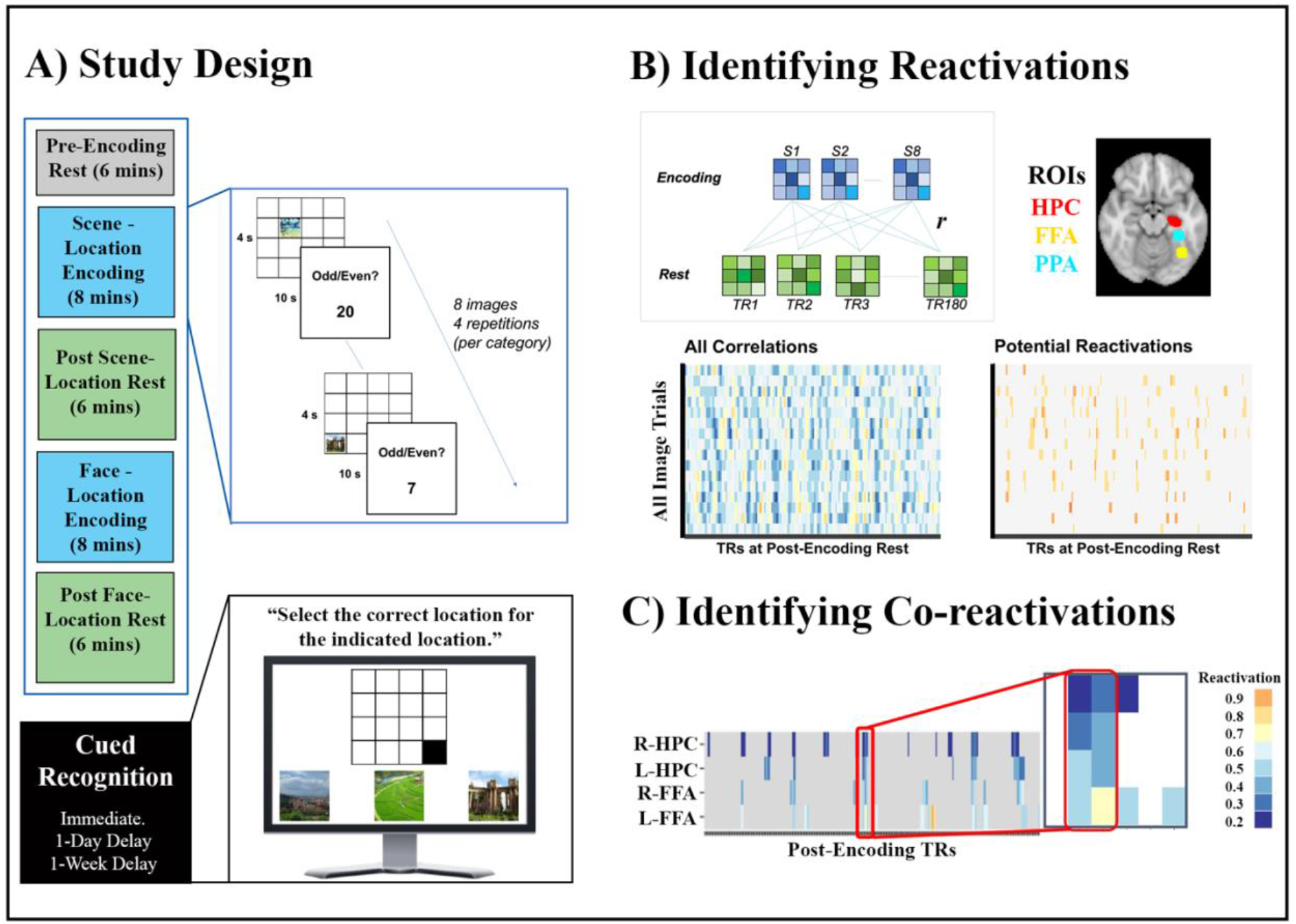
**A.** Study design. Participants learned image-location pairs in the scanner, and then completed a cued-recall task outside the scanner. **B.** Display of reactivations in HPC for one representative subject. Each image trial’s encoding neural patterns (beta weights) across all the voxels were correlated with the preprocessed pattern of activity across all voxels at each TR of the post-encoding rest period (separately conducted for HPC, FFA, and PPA). This results in a correlation matrix between all the image trials and all the rest period TRs, which is then thresholded to reflect “potential reactivations”. **C.** Illustration of example co-reactivating TRs (defined as reactivation in two ROIs to the same image trial, at the same TR (+/− 1 TR).

## Materials and Methods

### Participants

Thirty-six right-handed young adults (M_age_ = 22.1, SD = 3.16, 20 Females) from Temple University and the surrounding community participated in the study between 2019 and 2022. All participants spoke English and were free of neuropsychiatric disorders and MRI contraindications. Data collection was disrupted due to the Covid-19 pandemic, resulting in two cohorts of 21 participants and 15 participants, respectively. One participant from the first cohort, and two participants from the second cohort were excluded from the analyses due to little-to-no event-related MRI signal. Comparisons between the cohorts regarding their behavioral memory performances and imaging quality did not reveal any significant differences (see supplementary materials (SM) for Table S1), therefore we collapsed the data across the two cohorts for all the reported analyses, resulting in a sample of 34 participants.

### Study Design

Participants completed three rest and two task (encoding) runs (Fig. 1A). Scanning started with a baseline (*pre-encoding* henceforth) rest run (6 minutes (m)), which was used as a control rest period for reactivation and co-reactivation analyses (see below, *Co-reactivation Analysis*). Participants then completed two encoding task runs (8 m each), each of which was followed by a post-encoding rest period (6 m each). The order of encoding blocks (face-first or scene-first) were counterbalanced across participants. Total fMRI scan time was 34 m.

The task used an event-related design. Each trial began with a fixation cross (1 s) followed by a face or scene (4 s) located in a specific location on a four-by-four grid (‘image trial’). This was followed by a 10-s long odd-or-even number-judgement task during which time they were shown random numbers (2 s each) and were asked to press a button to indicate whether it was an odd or even number (referred to as ‘number trials’). The purpose of the number trials was to provide time to allow us to model the fMRI signal for each image trial while imposing a task that dampened overt rehearsal. Total trial length was 16 s. There was a total of eight images per face/scene category, and each image was repeated four times in total. Face and scene images were paired with different parts of the grid to avoid any overlap between the paired associates across categories, and their distribution across the four quadrants of the grid were also counterbalanced. Participants were instructed to pay attention to the image trials, to press a button on the number trials, and to keep their eyes open during the rest scans.

Upon exiting the scanner, memory was tested immediately, 1-day later and 1-week later, with two surprise tasks: a cued-location recall and a cued-image recognition task. Cued recall and recognition were completed in separate blocks, and their order was counter-balanced across participants. In this manuscript, we focus on the recognition task because the cued-location recall task showed susceptibility for “new learning” occurring during the repeated testing contexts, such that this measure did not show forgetting over delays. During recognition, participants were shown a four-by-four grid, with one of the cell locations highlighted with black, together with three image options (one target and two old lures from the same category which were also previously encoded). They were instructed to ‘choose (click on with the mouse) the correct image that had been paired with the highlighted location during encoding’. There were 16 recognition trials, evenly split between faces and scenes. A categorical accuracy variable (correct/incorrect) was derived for each trial, and a total correct variable was created across all the trials for each participant.

### Behavioral Analyses

Using separate Chi-square tests, we first tested the frequency of correct versus incorrect trials for recognition for each test day (Fig. 2). We then examined whether there were significant changes in trial-based recognition accuracy across the three test days. To this end, we first created three 2-by-2 contingency tables with the trial counts for behavior change from immediate to 1-day delay, from immediate to 1-week delay, and from 1-day to 1-week delay conditions (see *Table S2*). For these tables, behavior change was coded as “correct at both tests”, “initially correct – then incorrect”, “initially incorrect – then correct”, and “incorrect at both tests”. Since these are paired data, we then conducted three separate McNemar’s tests (McNemar, 1947) for the behavioral change across days.

**Fig 2.**
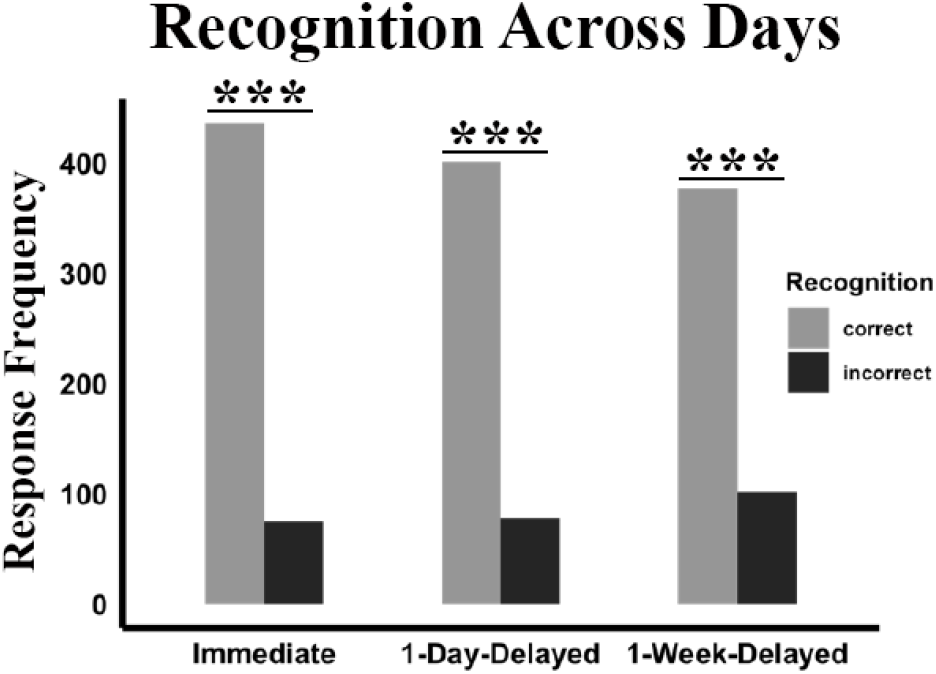
Distribution of correct and incorrect trials across the three tests. ***:*p* < 0.001

### fMRI Data Acquisition and Quality Check

MRI scans were completed at Temple University on a 3T Siemens Magnetom Prisma scanner, using a 64-channel phased-array head coil. High-resolution T1-weighted anatomical images were collected using a three-dimensional magnetization prepared rapid acquisition gradient-echo pulse sequence (TR = 520ms, TE = 0.007ms, FOV = 100 mm, flip angle = 60 ^o^, 2 mm slice thickness). Functional T2*-weighted images were collected using a gradient-echo planar pulse sequence with the following parameters: TR = 2000ms, TE = 29ms, FOV = 100 mm, flip angle = 76°, 2 mm slice thickness. DICOM images were converted to NIFTI format with Brain Imaging Data Structure (BIDS) nomenclature using dcm2niix (Gorgolevski et al. 2016). Quality control was achieved by running the MRIQC pipeline (version 0.10.4 in a Docker container) (Esteban et al. 2017) on the structural and functional images.

### fMRI Preprocessing

fMRI preprocessing was performed with FSL 6.0.1. (Jenkinson et al., 2012). First, the T1-weighted (T1w) anatomical image was skull stripped using the Brain Extraction Tool (BET). This image was used to assist in spatial normalization processes detailed below. Brain tissue segmentation of white matter (WM), gray matter (GM), and cerebrospinal fluid (CSF) was performed on the brain extracted T1w images using FAST. These segmentations were used to extract time series from the WM and CSF for reduction of noise in our preprocessing stream. FMRI preprocessing was completed using the fMRI Expert Analysis Tool (FEAT) version as implemented in FSL 6.0.1. using a pipeline designed to minimize the effects of head motion (Murty et al., 2018). This included simultaneous head motion correction, and non-linear warping to the MNI space, but no temporal or spatial filtering. The same preprocessing methods were applied to all encoding and rest runs.

### Defining Regions of Interests

Hippocampus (HPC), fusiform face area (FFA) and parahippocampal place area (PPA) were *a priori* selected for their role in memory and category selective processing, respectively, and were functionally defined in the MNI space at the group level. We first extracted univariate activity for each encoding task, and then completed a higher-level statistical analysis in fMRI Expert Analysis Tool (FEAT) version as implemented in FSL 6.0.1. to extract a group average of face>scene and scene>face contrasts. Active clusters from the face>scene contrast was used to extract the peak coordinates for FFA (MNI: left: x = 64, y = 43, z = 24; right: x = 23, y = 41, z = 25), while active clusters from the scene>face contrast was used to extract the peak coordinates for PPA (MNI: left: x = 57, y = 37, z = 31; right: x = 29, y = 44, z = 29), thresholding the z-stat maps at 3.1. Using a lower threshold (1.5), the peak hippocampal voxels that were active in both face>scene and scene>face contrasts were identified for HPC (MNI: left: x = 56, y = 47, z = 33; right: x = 34, y = 50, z = 32). For each ROI, we created a sphere using a 5mm radius kernel with the fslstats command in FSL around the peak coordinates. Importantly, each mask was created in MNI space and binarized before extracting any activity patterns from the task or rest scans.

### fMRI Multivariate Analysis

After preprocessing, we ran two separate general linear models (GLMs) for face-location and scene-location blocks, which modeled each image trial as a separate regressor. Importantly, each event regressor included all four repetitions of that image trial to increase detection power. Therefore, each GLM included 8 event regressors, each modeled for 4 seconds duration and were convolved with a double-gamma hemodynamic response function. Six head-motion parameters, and their first derivatives, and time series extracted from cerebrospinal fluid and white matter were added as covariates to the model to reduce noise. Voxel-vise encoding activity was extracted from the t-stat maps for each image trial, within each ROI. We chose to use t-statistics because doing so addresses the noise from highly variable voxels (Misaki et al., 2010; Dimsdale-Zucker and Ranganath, 2019).

All GLMs were run using FEAT version 6.0 as implemented in FSL 6.0.3. First-level face>baseline and scene>baseline contrasts were estimated in our regions-of-interest (ROIs), separately for each hemisphere (see next section for ROI selection). Finally, we modeled all three rest scans (i.e., the pre-encoding and two post-encoding rests) in GLMs with the same nuisance parameters. We then obtained and high-pass filtered the residuals from these models, and extracted TR-based activity from the residual t-stat maps.

### Coordinated Reactivation (Co-Reactivation) Analysis

First, reactivation of encoding events was quantified using representational similarity analysis (RSA; Kriegeskorte, Mur & Bandettini, 2008). Within each encoding session for each participant, each trial’s encoding patterns were correlated (Pearson’s correlation coefficients) to the pattern of activity at each TR of the rest periods to find potential reactivations. Importantly, we completed this analysis in both pre- and post-encoding rest periods, which resulted in a total of four (face-pre-encoding, face-post-encoding, scene-pre-encoding, and scene-post-encoding) 8 (encoding trials) by 180 (rest TRs) reactivation matrices (see Fig. 1B for an example) for each ROI. All correlations were then Fisher z-transformed.

Theoretically, there should not be any “reactivations” during pre-encoding rest, given that the participants did not see any of the image trials before. Thus, pre-encoding similarity patterns were used for thresholding the reactivation patterns that were detected at post-encoding: We first calculated the average similarity for each image trial across all pre-encoding-rest TRs, and then defined a 1.5 SD above the pre-encoding average threshold (1.5 value based on prior work by Staresina et al., 2013 and Schapiro et al., 2018). Only the post-encoding TRs that survived each image’s calculated threshold were considered potential ‘reactivations’ for that image trial (Fig. 1B). Next, we counted the number of such *reactivating* TRs to define our reactivation count variable. This analysis was repeated for all our ROIs, and separately using post-face rest for face-location trials, and post-scene rest for scene-location trials.

We next counted TRs (+/−1) that reactivated the same image trial across our ROIs. TRs that reactivated the same image trial across one of the HPCs (left or right) and one of the cortical regions (left or right FFA (or PPA)) were counted as *co-reactivating* TRs for two ROI pairs: HPC-FFA, and HPC-PPA (Fig. 1C). All reactivation and co-reactivation analyses were completed in a custom MATLAB code (version 2020b, available at https://www.mathworks.com/products/new_products/release2020b.html).

Importantly, our a priori hypothesis was that FFA should reactivate faces (Kanwisher and Yovel, 2006) and PPA should reactivate scenes (Epstein and Kanwisher, 1998; Park and Chun, 2009) given their suggested role in category-selective processing. However, we did not find any significant differences between FFA and PPA in selectively reactivating faces and scenes, respectively, therefore we collapsed trials across categories for all reactivation and co-reactivation analyses. This collapsed data included post-face rest for face trials, and post-scene rest for scene trials as their respective *post-encoding* rest in all the reported analyses.

### Testing the Relationship between Reactivation and Memory Performance

Using item-level multi-level linear models, entering the subject information as random slopes, we tested whether reactivation counts were predicted by rest period (post-encoding versus pre-encoding), separately for each ROI. The results were Bonferroni corrected for multiple comparisons at *p_adjusted_* = 0.008. For the ROIs that showed significantly higher reactivation at post-encoding than pre-encoding rest, we then tested whether these reactivations were associated with recognition accuracy (Fig. 3).

**Fig 3.**
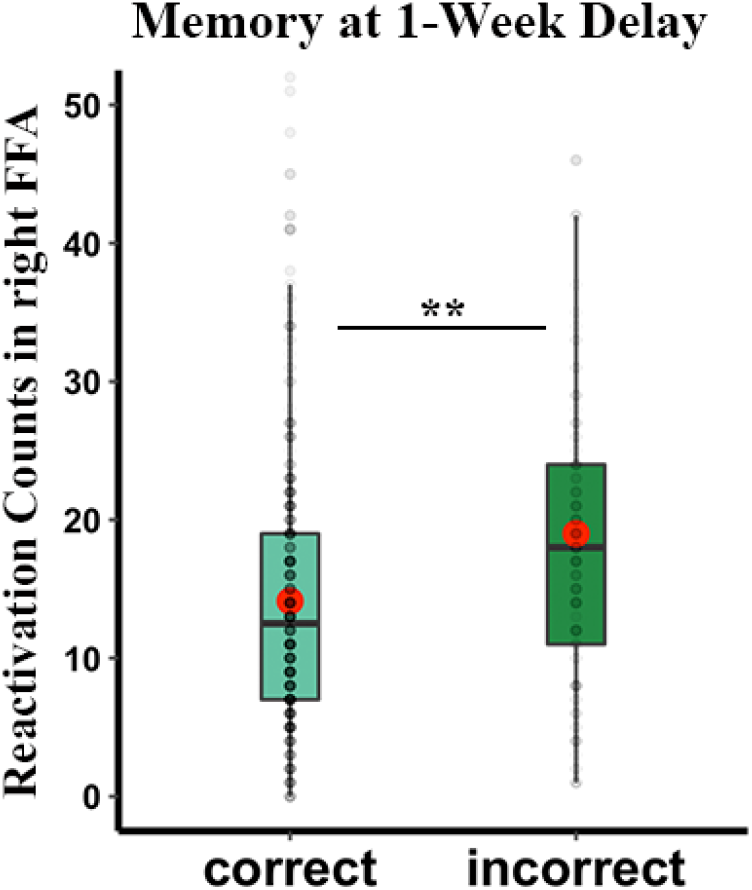
Incorrectly recognized image trials are associated with increased post-encoding reactivation counts in right FFA. Gray dots represent individual data points, i.e., reactivation count for a specific image trial. Red dots demonstrate the average reactivation counts. **: *p* < 0.01

### Testing the Relationship between Co-reactivation and Memory Performance

We next tested similar models to predict co-reactivation counts from the rest periods for each ROI pair (Bonferroni corrected at *p_adjusted_* = 0.0125). Using these results as a filter (e.g., retaining the ROI pairs that showed significantly higher co-reactivation at post-encoding than pre-encoding rest), we then tested whether the co-reactivation counts were significantly related to memory recognition (Fig. 4).

**Fig 4.**
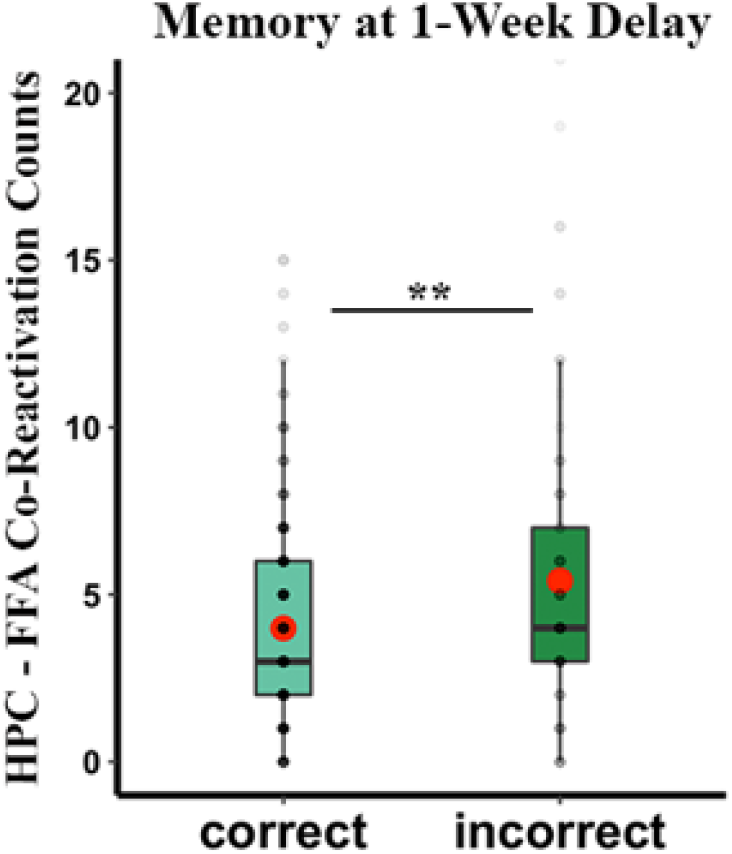
Incorrectly recognized image trials are associated with increased post-encoding co-reactivation counts between HPC and FFA. Gray dots represent individual data points, i.e., co-reactivation count for a specific image trial. Red dots demonstrate the average co-reactivation counts. **:*p* < 0.01

The unstandardized beta coefficients are reported for all our significant results. Reported statistical analyses were performed using R software (R package version 3.4.1) using mcnemar.test (rcompanion library), the cor.test, t.test, aov, lm (the stats library), and lmer (the lme4 library) functions depending on the test. All continuous variables were standardized before testing the regression models. Analysis scripts are available upon request.

## Results

### Recognition Accuracy Changes from Immediate to 1-Week-Delay Testing

Using separate Chi-square tests, we first tested the frequency of correct versus incorrect responses for recognition for each test day. These tests revealed that across all test days, participants had significantly higher numbers of correct than incorrect responses (Immediate: *X^2^*(1) = 256, *p* < 0.0001; 1-Day-Delay: *X^2^*(1) = 219, *p* < 0.0001; 1-Week-Delay: *X^2^(1)* = 159, *p* < 0.0001). Given that participants repeated the same recognition memory test, we next asked how their accuracy changed across days for image trials. Using McNemar’s Test, we found that there were no significant changes in behavior from immediate to 1-day delay condition (*p* = 0.2), or from 1-day to 1-week delay (*p* = 0.2) conditions. Importantly, however, participants’ responses indicated significant forgetting from the immediate to 1-week delay test (McNemar’s *X^2^*(1) = 9, *p* = 0.003).

### fMRI Event-Specific Reactivation During Post-Encoding Rest Period were Associated with Incorrect Recognition

Using representational similarity analysis (RSA), we first identified pre- and post-encoding TRs that showed significant reactivations of each image trial in our *a priori* ROIs (Fig. 1B). Our category-specific tests showed non-differentiated reactivation counts in FFA and PPA for faces and scenes, and we collapsed across categories for the following reactivation and co-reactivation analyses.

Using item-level multi-level linear modeling, we found that the hippocampus showed a greater number of detected “reactivations” during the post-compared to the pre-encoding rest period (Left: *F*(34,1024) = 2.5, *β* = 0.9, *SE* = 0.36, *p* = 0.013; Right: *F*(34,1024) = 4.72, *β* = 1.93, *SE* = 0.41, *p* < 0.001) suggesting that the defined reactivation patterns reflect experience-dependent changes of the encoded event. Similarly, the FFA showed more reactivations during the post-than pre-encoding rest, with these effects being more evident in the right hemisphere (Left: *F*(34,1024) = 1.87, *β* = 0.76, *SE* = 0.41, *p* = 0.062; Right: *F*(34,1024) = 6.19, *β* = 2.97, *SE* = 0.48, *p* < 0.001). Finally, PPA showed higher reactivations during post-than pre-encoding rest, although these effects were more prominent in left than right hemisphere (Left: *F*(34,1024) = 2.14, *β* = 0.79, *SE* = 0.37, *p* = 0.032; Right: *F*(34,1024) = 1.91, *β* = 0.64, *SE* = 0.33, *p* = 0.056). The effects in right HPC and right FFA survived Bonferroni correction (*p_adjusted_* = 0.008).

Next, we asked whether event-specific post-encoding reactivations were associated with subsequent recognition memory test performance (see *Methods* and *SI* for details). Interestingly, we found that there were a greater number of reactivations in right FFA for trials that were *incorrectly* recognized on the 1-week delayed recognition task compared to correctly recognized image trials (*F*(34,1024) = 2.9, *β* = 3.27, *SE* = 1.13, *p* = 0.004) (Fig. 3). Importantly, the number of “reactivations” during the pre-encoding rest period did not differ based on subsequent memory performance (*F*(34,1024) = −0.99, *β* = 0.49, *SE* = 0.5, *p* = 0.33), indicating that the phenomenon is experience-dependent. No other ROIs showed significant differences in reactivation counts based on subsequent recognition performance after a 1-week delay (see *SI* for more details).

### fMRI Event-Specific Co-Reactivation During Post-Encoding Rest Period were Associated with Incorrect Recognition

The prior analyses looked at one region in isolation, but as detailed above, to truly test more systems-like consolidation we wanted to assess coordinated reactivation across the hippocampus and cortex. Thus, we identified post-encoding TRs that showed significant co-reactivation to the same image trials in HPC and at least one cortical ROI (e.g., FFA or PPA, Fig. 1C). Using item-level multi-level linear modeling, we tested a model to predict the number of co-reactivation TRs (*co-reactivation counts* henceforth) from the post- and pre-encoding rest periods, separately for each of the hippocampal-cortical ROI pairs. We found that there were significantly higher co-reactivation counts for the HPC and FFA during the post-encoding compared to pre-encoding rest periods (*F*(34,1024) = 4.33, *β* = 0.79, *SE* = 0.18, *p* < 0.001). There was no difference in counts for HPC and PPA (*F*(34,1024) = 1.7, *β* = 0.3, *SE* = 0.17, *p* = 0.09), therefore we focused the remaining analyses on HPC-FFA co-reactivations.

We next tested whether the HPC-FFA co-reactivation counts during post-encoding rest were associated with recognition performance. This analysis revealed that there were higher HPC-FFA co-reactivations counts for incorrectly recognized images at a 1-week delay (*F*(32,480) = 2.89, *β* = 1.18, *SE* = 0.41, *p* = 0.004) (Fig. 4; see Fig. S3 for the immediate and 1-day delayed conditions), but not the earlier memory tests. Importantly, there was no significant difference in subsequent memory for items as a function of “co-reactivation” during the pre-encoding rest period (*p* = 0.67), further suggesting that the effects observed at post-encoding rest are experience dependent.

The above analyses, showing a negative relationship between co-reactivations and memory at a 1-week delay, suggests that these co-reactivation signals occurring after rest reflect consolidation-related process that might actually impair associative recognition. Importantly, we did not show this significant inverse relationship for the immediate and 1-day delayed tests, further implicating a role in consolidation. A more specific way to test whether this relationship is related to consolidation is to use a measure of forgetting, in which we compare memory across testing delays. Accordingly, we coded image based on their relative status at the immediate and delayed tests. Image trials that were correct at immediate and remained correct at the 1-day or 1-week delayed test as “remembered”. Image trials that were initially correct at immediate but were incorrect at the 1-day or 1-week delayed test were coded as “forgotten”. We then tested whether there was any significant relationship between HPC-FFA co-reactivations and remembered and forgotten trials, tested separately for changes from immediate-to-1-day delayed, and immediate-to-1-week delayed. We found a consistent pattern of relationship for forgetting after 1-day and forgetting after a week. Specifically, we found that trials that were initially correct at immediate but then were forgotten at 1-day-delay showed a greater number of post-encoding HPC-FFA co-reactivations (*F*(32,414) = 2.03, *β* = 1.09, *SE* = 0.54, *p* = 0.043). Similarly, we found that trials that were initially correct at immediate but then were forgotten at the 1-week-delay condition showed a greater number of post-encoding HPC-FFA co-reactivations (*F*(32,407) = 2.16, *β* = 1.02, *SE* = 0.47, *p* = 0.031) (Fig 5). We did not find any significant differences in the number of HPC-FFA co-reactivations for remembered and forgotten trials when we tested the model for changes from 1-day to 1-week delay testing (*F*(31,386) = 1.58, *β* = 0.78, *SE* = 0.5, *p* = 0.11). Together, these findings suggest that post-encoding HPC-FFA co-reactivations were uniquely associated with *forgetting* over time.

**Fig 5.**
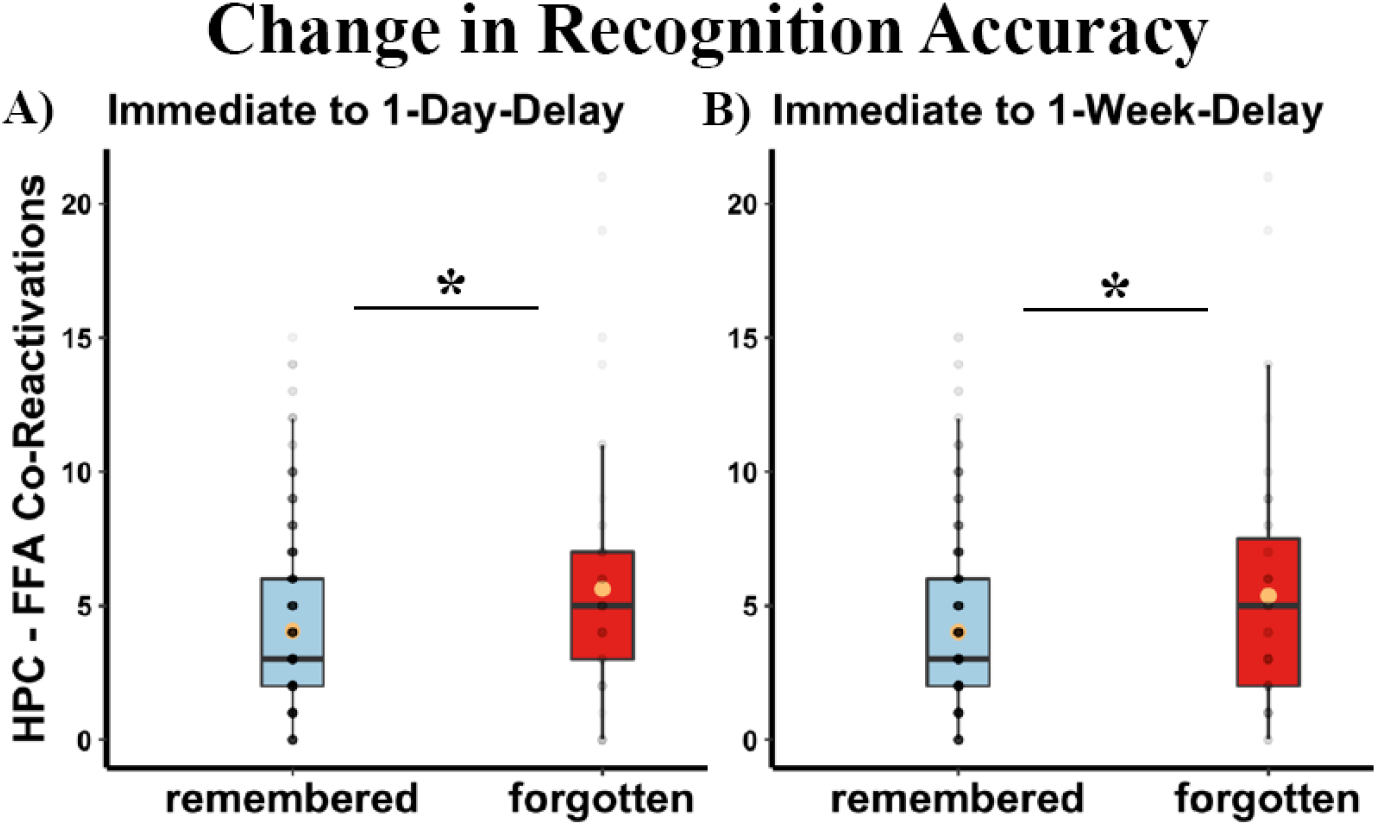
Images that were forgotten in time showed greater number of HPC-FFA co-reactivations at post-encoding rest. **A.** Recognition accuracy change from immediate to 1-day delay condition (*p* = 0.043). **B.** Recognition accuracy change from immediate to 1-day delay condition (*p* = 0.031). *Remembered:* image trials that were correct at immediate testing and remained correct in the respective subsequent test. *Forgotten:* image trials were correct at immediate testing but were then incorrect in the respective subsequent test. Yellow dots represent the average co-reactivation counts. *: *p* < 0.05

### Specificity of Co-Reactivation Effects on Recognition Memory

Given our initial observation that greater reactivation counts in right FFA were significantly associated with incorrectly recognized image trials at 1-week delay, we asked whether the association between HPC-FFA co-reactivations and incorrect delayed recognition performance was driven by the reactivations in right FFA alone. To this end, we examined TRs in right FFA that co-reactivated a given image with HPC, which we refer to as ‘coordinated’, versus right FFA TRs that did not show any co-reactivations (‘uncoordinated’). Using the total number of coordinated and uncoordinated TRs in right FFA, we then re-tested our original model for the right FFA and 1-week-delayed recognition association. The results revealed that the observed relationship was specific to *coordinated* reactivations. Namely, the number of coordinated TRs, but not uncoordinated TRs, in the right FFA was greater for incorrectly recognized compared to correctly recognized trials at 1-week-delay (coordinated: *F*(32,480) = 2.48, *β* = 1.16, *SE* = 0.47, *p* = 0.014, uncoordinated: *F*(32,480) = 1.82, *β* = 1.21, *SE* = 0.67, *p* = 0.07) (Fig. 6). Further, we did not find any significant relationship between 1-week delayed-recognition and uncoordinated TRs in HPC (uncoordinated in left HPC: *F*(32,480) = 0.16, *β* = 0.08, *SE* = 0.53, *p* = 0.88; uncoordinated in right HPC: *F*(32,480) = 1.36, *β* = 0.76, *SE* = 0.56, *p* = 0.18). Together, these findings support the conclusion that it is the coordination of the HPC and FFA reactivations that is predictive of poorer subsequent recognition, rather than merely the reactivation of an individual region. In other words, when these regions reactivate independently, they don’t show any significant relationship with subsequent memory. But when they co-reactivate, they are associated with forgetting.

**Fig 6.**
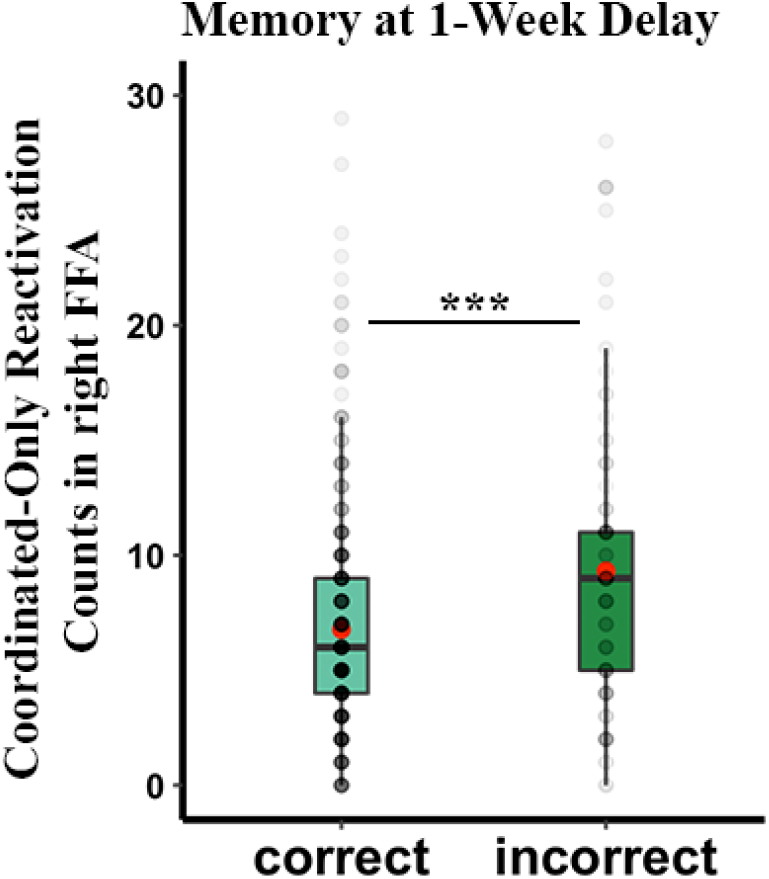
Coordinated-Only Reactivations in Right-FFA. Coordinated TRs in right FFA were associated with incorrect recognition. Gray dots represent individual data points; red dot demonstrates the average reactivation counts. ***:*p* < 0.005.

## Discussion

For the last two decades, researchers have shown that in rodents, hippocampal reactivations during sleep or awake rest recapitulated previous events, and in some cases, correlated with better subsequent memory (Buzsáki, 1989; Wilson and McNaughton, 1994; Skaggs and McNaughton, 1996; Foster and Wilson, 2006; Diba and Buzsáki, 2007; Davidson et al., 2009; Karlsson and Frank, 2009; Girerdeau and Zugaro, 2011; Jadhav et al., 2012). More recently, this work has been extended to show that the hippocampus and cortex can also co-replay previously encountered locations (Ji and Wilson, 2007; Lansink et al., 2009; Wierzynski et al., 2009; Peyrache et al., 2009).

Here, we asked whether in humans, a similar sort of co-replay of individual events occurs, and how it affects subsequent memory. First, using a novel method (TRCR), we showed that there was an increased number of co-reactivations between the HPC and fusiform face area (FFA) for individual experienced events during post-encoding (as compared to pre-encoding) awake rest. Moreover, not only was the frequency of co-reactivations significantly higher at post-than pre-encoding, but also the strength of these co-reactivations (see supplementary materials). Importantly, the appearance of increased HPC-FFA co-reactivations during the post-encoding rest period strongly indicates that these co-reactivation patterns reflect experience-dependent changes related to encoded events, and not just spurious similarities. Second, we found that HPC-FFA co-reactivations were associated with diminished memory performance at 1-week delay, e.g., corresponded with the increase in incorrectly recognized trials. Finally, we found that these co-reactivations were specifically associated with the forgetting of trials that had been initially correctly recognized at immediate testing, suggesting that the observed phenomenon is not merely the result of failed encoding, but rather, is linked to post-encoding consolidation processes that predict the later weakening or blurring of representations.

### Systems Consolidation and Coordinated Reactivation

Some prior fMRI work has shown that item-specific post-encoding reactivations were associated with better subsequent memory (e.g., Deuker et al., 2013; Staresina et al., 2013). As such, the seemingly paradoxical findings from the present evaluation of how *co*-reactivations relate to memory raise important questions about exactly how hippocampal-cortical co-reactivations may transform event memories in addition to, or differently from, regionally limited reactivations as they have been more traditionally assessed.

Systems consolidation theories posit that hippocampal and cortical representations are different in nature: the hippocampus is posited to be engaged in pattern separation (Yassa and Stark, 2011), supporting storage of more detailed episodic representations, whereas the cortex is theorized to generalize and store a more gist level representations of events (Kumaran et al., 2016; Moscovitch and Gilboa, 2022). Importantly, these different representations can co-exist and interact with one another, depending on the task demands (Winocur and Moscovitch, 2011; Sekeres et al., 2018; Moscovitch and Gilboa, 2022). This interaction is likely reflected in hippocampal-cortical co-reactivations. However, any expected impacts of co-reactivations on subsequent memory are not fully fleshed out in current memory consolidation theories. One possibility is that these different representations could bias the memory decision in opposing directions: while cortical representations might inform gist-dependent memory judgments, hippocampal representations might bias memory judgment towards unique details. Any conflict between the gist representation with the specific event representation could lead to interference, hence forgetting. We speculate that our central observation – that co-reactivations relate to incorrect recognition and forgetting – may be explained by conflicts that arise when the hippocampus and cortex attempt to represent and communicate different versions of events; with hippocampal reactivations pushing toward pattern separation among image representations, but cortical reactivations evincing more gist-like memory representations.

Another possibility regarding the interaction of co-existing hippocampal and cortical representations is that it reflects cross-regional interactions that lead to the integration of related events, and that this integration itself may disrupt subsequent representation of event-specific encoding patterns. This interpretation would be more in line with trace transformation theories of consolidation, which proposed that memories undergo re-organization during consolidation, through which they are transformed into more generalized variants of themselves, retaining gist but losing details and context-specificity (Sekeres et al., 2018; Moscovitch and Gilboa, 2022). Consistent with this idea, Tompary and Davachi (2017) showed that events that share overlapping associations have become increasingly similar in their cortical (in this case mPFC) representations over time. Moreover, this increased similarity for overlapping events was associated with increased hippocampus-mPFC functional connectivity during post-encoding rest, providing further evidence that post-encoding processes help prioritize integration of related events through their commonalities. As stated above, this prioritization of common features during consolidation processes may result in forgetting of unique features that would otherwise enable recollection of individual events. This account provides an alternative interpretation for why we find higher HPC-FFA co-reactivations for trials that are incorrectly recognized at 1-week delay. It is likely that these co-reactivations have blurred out the unique features of these events, thereby making it difficult to distinguish the target image from the two other alternative lure images, which were themselves also studied during encoding. Indeed, using a small number of grid locations as the paired associates to these images may have biased the task toward producing such generalization, especially since the grid locations on their own do not provide any discriminable features that could help to uniquely identify individual events. This state of affairs might be different in important ways from previous tasks in which reactivated items were paired with common associates of the items (e.g., pairing a cat image with a meow sound; Oudiette et al., 2013).

We did not find any significant relationship between HPC-FFA co-reactivations and recognition at immediate or 1-day delay testing. The fact that these effects have only emerged after a 1-week delay is in line with the consolidation hypotheses that memories are transformed over time (Kumaran et al., 2016; Moscovitch and Gilboa, 2022).

### Reactivation may have different effects in different states of consciousness

So far, we have offered several alternative interpretations, aligning with different assumptions from systems consolidation theories, for our finding that HPC-FFA co-reactivations are associated with incorrect recognition. However, it is also possible that the mechanisms underlying our findings may be better explained by theoretical frameworks distinct from consolidation theory. Here, we consider an important question regarding the function of awake (co-)reactivations: Does awake spontaneous post-encoding reactivation always benefit memory?

Awake reactivations have been considered highly similar to reactivations during sleep (Tambini and Davachi, 2019), yet it is likely that reactivation processes, and their outcome, may differ across different mental states. One study provided early evidence that awake rest may not be as protective against interference as previously thought. Diekelmann and colleagues (2011) compared awake versus sleep groups for the effect of cued reactivation on performance in an interference learning paradigm. Each group first learned object-location information, then underwent cued reactivation (through odor vs vehicle), and finally completed a new location learning task (interference). The authors found that cued reactivation during sleep was associated with improved object-location memory after interference learning but, in the wake group it was associated with *reduced* object-location memory due to interference learning. This finding, in line with other reconsolidation studies, suggests that awake reactivations may make memories more labile, allowing changes or updates to memory (Schiller et al., 2010; Kuhl et al., 2010; see Jardine et al., 2022 for a review), but also creating opportunities for forgetting due to interference (Diekelmann et al., 2011). Another study from the same group later found that cued reactivation during awake rest benefits only the cued items, whereas benefits of cued reactivation is extended to uncued items from the same context during sleep (Oudiette et al., 2013), further supporting the notion that reactivation during wakeful and sleeping periods may have different effects on memory. It is important to note that other dimensions of one’s mental state, beyond just awake versus asleep periods, may have substantive impacts on the functions of reactivation. Others have shown, for instance, reactivation differences in low versus high reward conditions (e.g., Gruber et al., 2016), and in threatening versus safe contexts (de Voogd et al., 2016). Together with these studies, our findings highlight the need for a more systematic investigation of the functions of reactivation as they arise in association with various mental states in humans.

### Limitations

The literature on memory reactivations in humans is very small, but is surprisingly varied with respect to methodology, because the laboratories conducting this type of research generally employ disparate tasks, stimuli, and memory testing methods (see Table 1). These methodological choices could potentially give rise to different results. In the spirit of transparency, we outline features of our task that may have given rise to our results. First, using a grid for associative learning may have contributed to an increased level of interference, due to the relatively low discriminability of grid locations (compared to using close associates of the presented items). Most prior fMRI work on reactivation has focused on pairs of visual items to test associative memory (e.g., Tambini et al., 2010). Notably, the Diekelmann et al (2011) study discussed above also utilized grid learning to specifically test how reactivations relate to later interference learning, and that study produced findings consistent with the present study with respect to wakeful reactivation, further supporting the idea that location-based learning may be particularly interference-inducing.

**Table 1.**
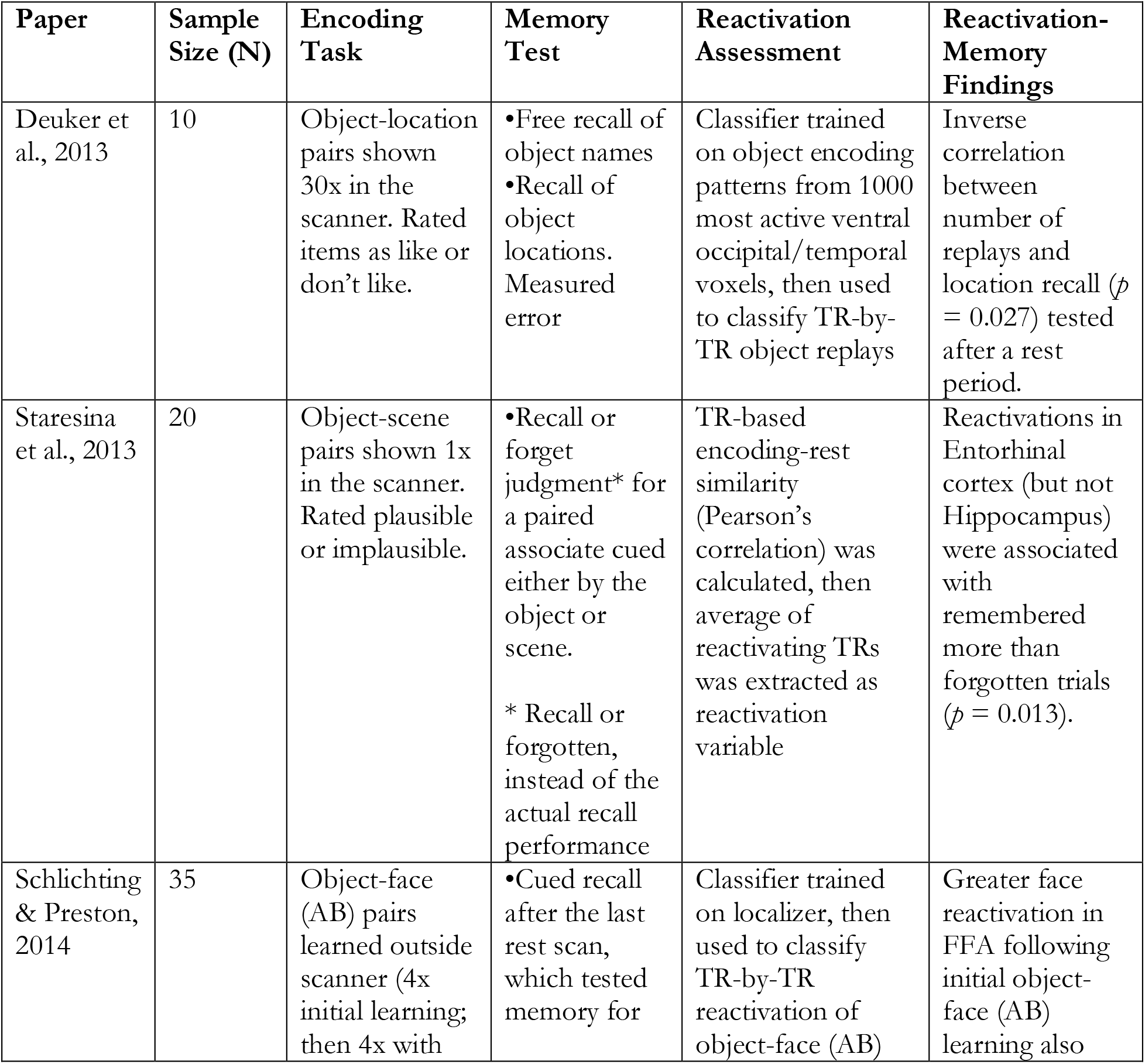
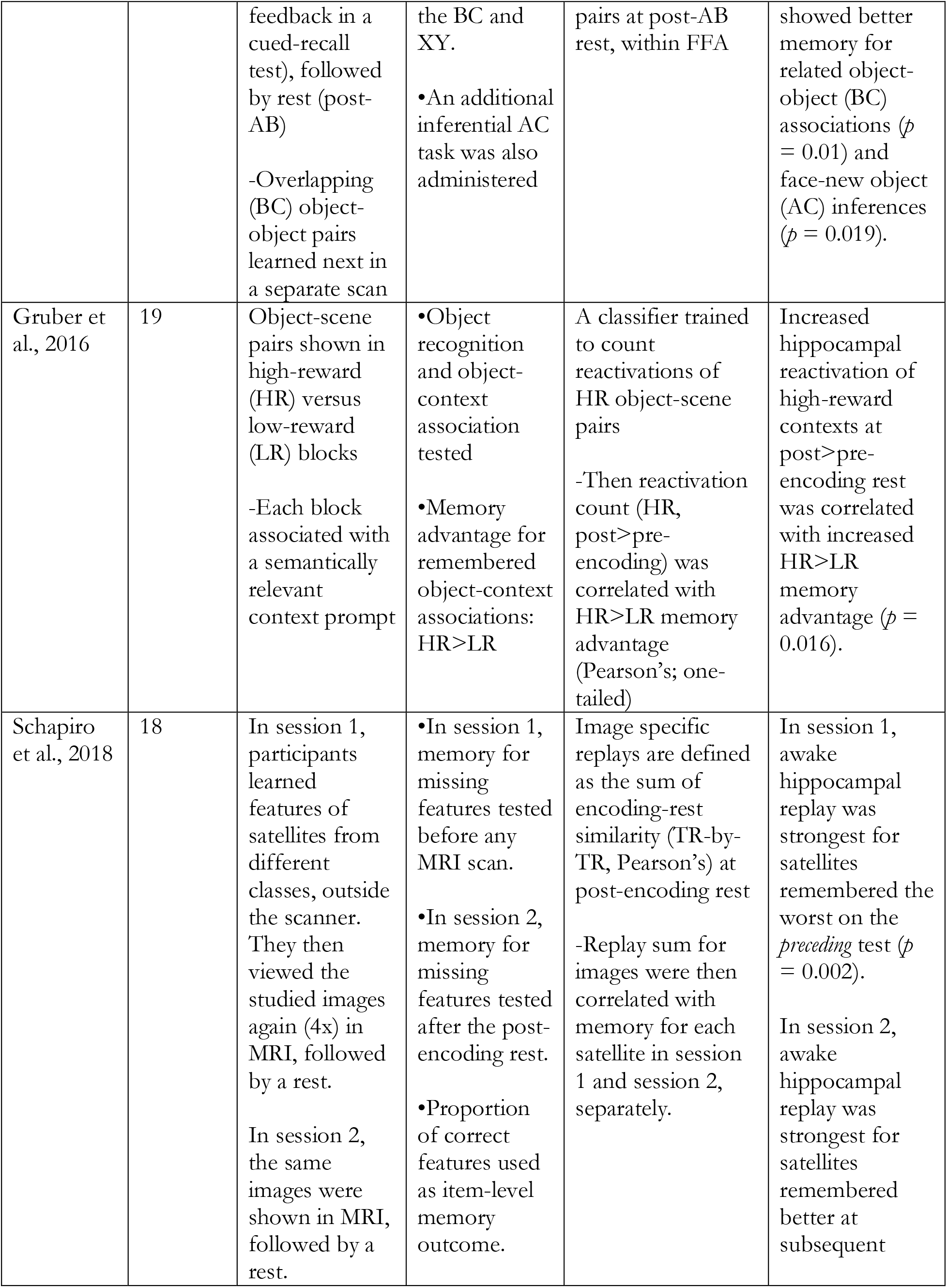
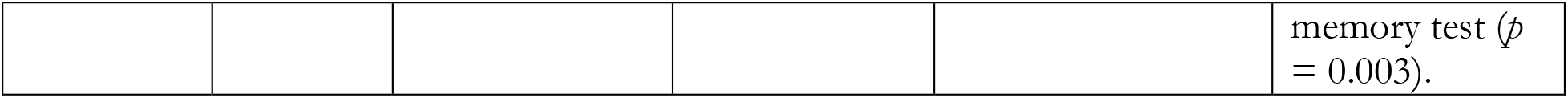
Summary of Previous Studies Investigating Event-Specific Spontaneous Reactivation at Awake Rest. Sample size is the final sample included in the analyses.

| Paper | Sample Size (N) | Encoding Task | Memory Test | Reactivation Assessment | Reactivation-Memory Findings |
| --- | --- | --- | --- | --- | --- |
| Deuker et al., 2013 | 10 | Object-location pairs shown 30x in the scanner. Rated items as like or don't like. | •Free recall of object names<br>•Recall of object locations.<br>Measured error | Classifier trained on object encoding patterns from 1000 most active ventral occipital/temporal voxels, then used to classify TR-by-TR object replays | Inverse correlation between number of replays and location recall ($p = 0.027$) tested after a rest period. |
| Staresina et al., 2013 | 20 | Object-scene pairs shown 1x in the scanner. Rated plausible or implausible. | •Recall or forget judgment* for a paired associate cued either by the object or scene.<br>* Recall or forgotten, instead of the actual recall performance | TR-based encoding-rest similarity (Pearson's correlation) was calculated, then average of reactivating TRs was extracted as reactivation variable | Reactivations in Entorhinal cortex (but not Hippocampus) were associated with remembered more than forgotten trials ($p = 0.013$). |
| Schlichting & Preston, 2014 | 35 | Object-face (AB) pairs learned outside scanner (4x initial learning; then 4x with | •Cued recall after the last rest scan, which tested memory for | Classifier trained on localizer, then used to classify TR-by-TR reactivation of object-face (AB) | Greater face reactivation in FFA following initial object-face (AB) learning also |
|  |  | feedback in a cued-recall test), followed by rest (post-AB)<br>-Overlapping (BC) object-object pairs learned next in a separate scan | the BC and XY.<br>•An additional inferential AC task was also administered | pairs at post-AB rest, within FFA | showed better memory for related object-object (BC) associations ($p = 0.01$) and face-new object (AC) inferences ($p = 0.019$). |
| Gruber et al., 2016 | 19 | Object-scene pairs shown in high-reward (HR) versus low-reward (LR) blocks<br>-Each block associated with a semantically relevant context prompt | •Object recognition and object-context association tested<br>•Memory advantage for remembered object-context associations: HR>LR | A classifier trained to count reactivations of HR object-scene pairs<br>-Then reactivation count (HR, post>pre-encoding) was correlated with HR>LR memory advantage (Pearson's; one-tailed) | Increased hippocampal reactivation of high-reward contexts at post>pre-encoding rest was correlated with increased HR>LR memory advantage ($p = 0.016$). |
| Schapiro et al., 2018 | 18 | In session 1, participants learned features of satellites from different classes, outside the scanner. They then viewed the studied images again (4x) in MRI, followed by a rest.<br>In session 2, the same images were shown in MRI, followed by a rest. | •In session 1, memory for missing features tested before any MRI scan.<br>•In session 2, memory for missing features tested after the post-encoding rest.<br>•Proportion of correct features used as item-level memory outcome. | Image specific replays are defined as the sum of encoding-rest similarity (TR-by-TR, Pearson's) at post-encoding rest<br>-Replay sum for images were then correlated with memory for each satellite in session 1 and session 2, separately. | In session 1, awake hippocampal replay was strongest for satellites remembered the worst on the *preceding* test ($p = 0.002$).<br>In session 2, awake hippocampal replay was strongest for satellites remembered better at subsequent memory test ($p = 0.003$). |

Second, our stimuli were repeated several times during encoding. This repeated presentation may have altered the encoding patterns, or their strength, thereby leading to differences in their (co-) reactivation. We think this variable is probably not essential to the pattern of our findings, given that previous work that has shown positive effects of reactivation on memory using varying numbers of repetition, from single shot learning (e.g., Staresina et al., 2013) to 30 repetitions (e.g., Deuker et al., 2013).

Third, our study tested recognition memory three times for all the learned image-location pairs. We acknowledge that this may have contributed to the current results in two different ways: On the one hand, as our behavioral findings suggest, there is a strong overall accuracy even after a 1-week delay, leaving us with limited behavioral variance across remembered and forgotten trials, which may be biasing the current results. On the other hand, it is possible that multiple testing sessions introduced additional interference over time, thereby leading to the forgetting that we observed over time.

Future research is warranted to systematically test whether or how any of these factors could have contributed to these findings. To that end, it is important to highlight that the current consolidation theories are, in general, critically lacking in the specification of boundary conditions: There is a lack of specificity regarding not only the types of (reactivation-like) processes that should occur, and how they should occur, but also *in which conditions* these processes are most likely to update, improve or weaken memories. Table 1 summarizes the different features of the previous work showing memory outcomes related with event-specific spontaneous reactivations at awake rest. We believe some of these methodological differences highlighted in Table 1 is a reflection of how the vaguely defined theoretical concepts are translated into different approaches to empirical design and analysis. Thus, we conclude that future research should systematically test how alternative design approaches impact reactivation effects on memory, and help define the boundary conditions for when and how reactivation, and co-reactivation, may improve or weaken event memories.

## Data Availability

Data and analysis scripts are available upon request to the corresponding author.

## Author Contribution

NM conceptualized and designed the study, KRJ collected the data. BT, ETC, VPM, JC, and IRO conceptualized the analyses. BT collected, analyzed, and visualized the data, and wrote the original draft. ETC and VPM engaged in writing, review and editing of the manuscript. VP, JC and IRO provided funding, supervision, and reviewed and edited the manuscript.

## Acknowledgements

We would like to thank Zachary S. Heffernan for their contributions during data collection.

