## SupplementaryMaterials for "Awake Hippocampal-Cortical Co-reactivation Is Associated with Forgetting"

**Supplementary Materials**

*Cohort Comparisons*

As reported, data were collected in two separate cohort. The two cohorts did not differ in age ($M_{difference}$ = 1.2, $p$ = 0.2), memory recognition performance ($M_{difference}$ = 0.01, $p$ = 0.9), or MRI signal quality checks (Fig. S1). Given the two cohorts' similarity in these metrics, their data were collapsed into one sample. Table S1 below reports the age, gender, race, income, and education for this collapsed sample.

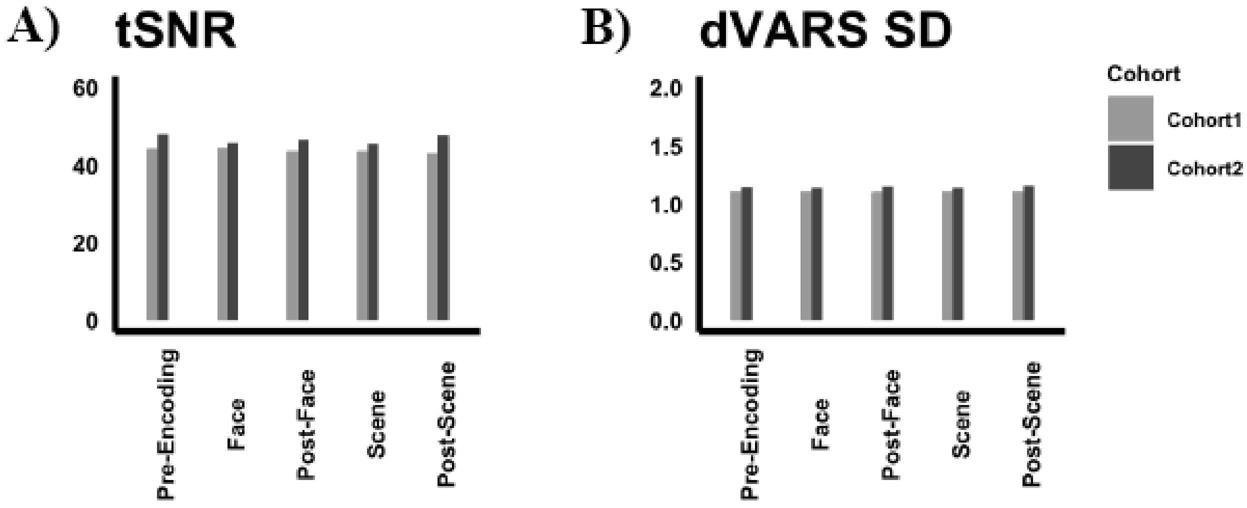


**Fig. S1. Cohorts did not differ in quality assurance metrics.** Differences in temporal signal-to-noise ratio (tSNR), standardized global signal (dVARS SD) were calculated for each functional run for both cohorts.

**Table S1. Demographics, Memory Recall Performance, and MRI Signal Quality**

| Characteristics | Mean (SD) or n (%) |
|---|---|
| Age, Years | 22.1 (3.16) |
| Gender, Female/Male | 20 (59%), 14 (41%) |
| Race | |
| White | 26 (77%) |
| Black | 1 (3%) |
| Asian, or Asian American | 2 (6%) |
| Mixed (Black, White) | 2 (6%) |
| Other | 2 (6%) |
| Family Income | |
| $ 15,000 or less | 3 (9%) |
| Between $15,001 and $34,999 | 4 (12%) |
| Between $35,000 and $49,999 | 3 (9%) |
| Between $50,000 and $74,999 | 0 (0%) |
| Between $75,000 and $99,999 | 2 (6%) |
| Greater than $100,000 | 5 (15%) |
| Not reported | 12 (35%) |
| Highest Education Completed | |
| Some college (at least 1-year) | 7 (21%) |
| Associate Degree | 2 (6%) |
| Bachelor's Degree | 6 (18%) |
| Some Graduate Training (degree not completed) | 3 (9%) |
| Graduate Degree (non-specified) | 1 (3%) |
| Not reported | 18 (53%) |

**Table S2. Memory Change: Contingency tables across days**

| | 1-Day Delay | | 1-Week Delay | |
|---|---|---|---|---|
| **Immediate** | **Correct** | **Incorrect** | **Correct** | **Incorrect** |
| **Correct** | 369 | 45 | 348 | 59 |
| **Incorrect** | 33 | 33 | 30 | 43 |

| | 1-Week Delay | |
|---|---|---|
| **1-Day Delay** | **Correct** | **Incorrect** |
| **Correct** | 331 | 55 |
| **Incorrect** | 40 | 38 |

*Post Encoding-Reactivation Count and Immediate and 1-Day Delayed Recognition Memory*

We tested associations between the post-encoding reactivation counts and 1-week-delayed recognition memory for all the ROIs that showed higher reactivations at post- than pre-encoding rest periods (left and right HPC, and left PPA). However, none of these regions showed any significant associations with the recognition memory at 1-week-delayed testing: right HPC ($F(32,480) = 1.51$, $\beta = 1.38$, $SE = 0.91$, $p = 0.13$), left HPC ($F(32,480) = 0.34$, $\beta = 0.26$, $SE = 0.78$, $p = 0.74$), or left PPA ($F(32,480) = 0.36$, $\beta = 0.3$, $SE = 0.86$, $p = 0.72$). Additionally, we tested whether post-encoding reactivations in right FFA were associated with recognition accuracy at immediate ($F(34,512) = 1.56$, $\beta = 2.04$, $SE = 1.3$, $p = 0.12$) and 1-day delayed testing ($F(34,512) = 1.01$, $\beta = 1.29$, $SE = 1.28$, $p = 0.31$). The results did not reveal any significant associations, but the effects were in the same direction, suggesting that the reactivation effects may become more prominent after a longer-delay than what is generally tested in similar reactivation research (Fig S2).

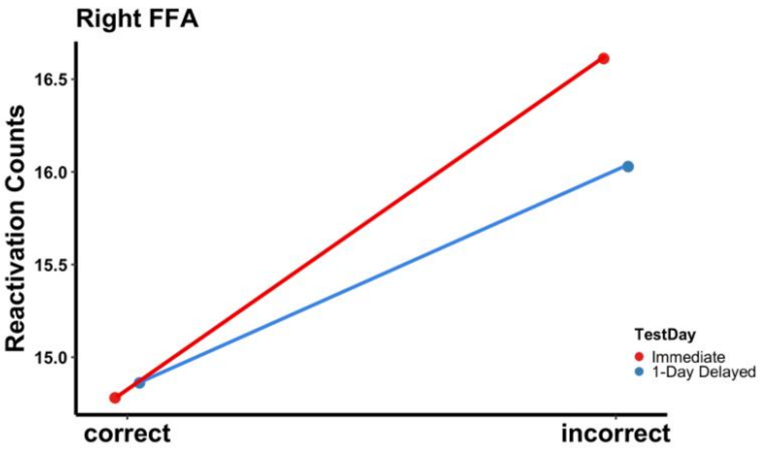


**Fig. S2. Post-Encoding Reactivation Count in right FFA was not significantly associated with A.** immediate (*p* = 0.12) or **B.** one-day delayed (*p* = 0.31) recognition accuracy.

*Post Encoding Co-Reactivation Count and Immediate and 1-Day Delayed Recognition Memory*

Similar to our reactivation analyses, we tested whether post-encoding HPC-FFA co-reactivations were associated with recognition accuracy at immediate ($F(34,512) = 1.0$, $\beta = 0.47$, $SE = 0.47$, $p = 0.32$) and 1-day delayed testing ($F(34,512) = 1.02$, $\beta = 0.46$, $SE = 0.45$, $p = 0.31$). The results did not reveal any significant associations, but the effects were in the same direction, suggesting that the reactivation effects may become more prominent after a longer-delay than what is generally tested in similar reactivation research (Fig S3). Additionally, given the trending level difference between HPC-PPA co-reactivations at post- and pre-encoding rest ($p = 0.09$), we tested whether the post-encoding HPC-PPA co-reactivations were associated with 1-week-delayed recognition memory. Our results revealed no significant associations ($F(32,480) = -1.06$, $\beta = -0.4$, $SE = 0.37$, $p = 0.29$).

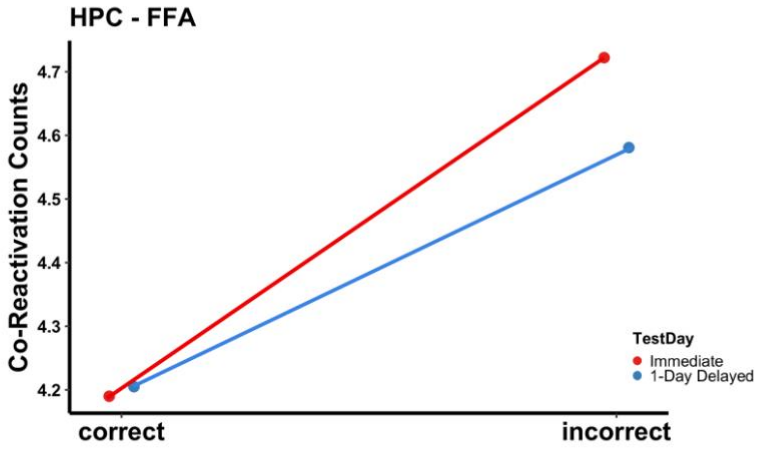


**Fig. S3. HPC-FFA Co-Reactivations** was not associated with recognition accuracy at immediate ($p$ = 0.32) and 1-day-delayed ($p$ = 0.31) testing**.**

*fMRI Event-Specific (Co-)Reactivation Sum Predicts Forgetting*

Our analyses focused on the total number of reactivations during post-encoding rest, but previous work has utilized sum (Schapiro et al., 2018) or mean (Staresina et al., 2013) of all awake reactivations as their "replay" variable. Thus, we asked whether our unique forgetting effect of awake reactivations is related to the variable of choice, that is reactivation counts. To test this, we derived a sum of reactivation scores for each unique event, i.e., calculated the sum of reactivation value (Pearson's $r$) across all the reactivation TRs. We then tested the same models reported in the main manuscript using this event-specific reactivation sum score. We found that hippocampus showed greater *sum* of reactivations during post- than pre-encoding rest period (Left: $F(34,1024) = 2.6$, $\beta = 0.31$, $SE = 0.12$, $p = 0.011$; Right: $F(34,1024) = 4.9$, $\beta = 0.7$, $SE = 0.14$, $p < 0.001$) suggesting that the defined reactivation patterns reflect experience-dependent changes in the image representations. Left FFA did not show any significant differences between pre- and post-encoding rest periods for the sum of reactivations more reactivations ($p = 0.47$), but right FFA had significantly higher reactivation sum during post- than pre-encoding rest ($F(34,1024) = 5.68$, $\beta = 1.26$, $SE = 0.22$, $p < 0.001$). Finally, PPA showed relatively higher reactivation sum during post-than pre-encoding rest, although these effects did not reach significance (Left: $F(34,1024) = 1.82$, $\beta = 0.28$, $SE = 0.15$, $p = 0.07$; Right: $F(34,1024) = 1.86$, $\beta = 0.29$, $SE = 0.17$, $p = 0.06$).

Next, we asked whether these event-specific post-encoding reactivation sums are associated with recognition performance in these regions showing this heightened reactivation sums (left, right HPC, and right FFA). Similar to our original results, we did not find any significant reactivation sum and recognition memory effects in neither left ($p = 0.78$) or right HPC ($p = 0.15$). We did replicate our effect in right FFA ($F(34,1024) = 2.98$, $\beta = 1.47$, $SE = 4.94$, $p = 0.003$) for incorrectly recognized image trials at 1-week delayed recognition. Importantly, this heightened reactivation *sum* – incorrect recognition association was not significant in pre-encoding rest period ($p = 0.34$).

We next tested HPC-cortex co-reactivations using the sum of co-reactivation (over all the co-reactivating TRs for a given pair). Replicating our findings with co-reactivation *counts*, we found significantly heightened sum of HPC-FFA co-reactivations at post- than pre-encoding rest ($F(34,1024) = 4.2$, $\beta = 0.61$, $SE = 1.46$, $p < 0.0001$). Moreover, this increased HPC-FFA co-reactivation sum was also associated with incorrect recognition at our 1-week-delayed test condition ($F(32,480) = 2.64$, $\beta = 0.86$, $SE = 0.32$, $p = 0.008$). Importantly, there was no significant HPC-FFA co-reactivation sum and recognition relationship at pre-encoding rest ($p = 0.68$). Finally, we did not show any significant differences between post- and pre-encoding rest regarding the HPC-FFA co-reactivation *sum* ($p = 0.1$), thereby eliminating any follow-up test for memory effects. Together, we replicated all our significant effects that relied on reactivation counts by using reactivation *sums*, suggesting that the difference between our findings and that of previous work (e.g., Staresina et al., 2013; Schapiro et al., 2018) cannot be attributed to the variable choice.

*fMRI Category-Level Reactivations Predict Forgetting*

Separate from our event-specific reactivation analyses, we also conducted reactivation analyses at the category-level. To this end, we first extracted the average of all images within a given category (face and scene), we then tested whether the patterns of these average face and scene images to reactivated and co-reactivated during the post-encoding rest period. Our analyses revealed that both right HPC ($F(34,128) = 3.33$, $\beta = 3.16$, $SE = 0.95$, $p = 0.001$) and right FFA ($F(34,128) = 2.59$, $\beta = 3.39$, $SE = 1.31$, $p = 0.01$) showed significantly higher number of reactivations during post- than pre-encoding rest. No other ROIs showed significant differences in the number of category-level reactivations (Left HPC: $p = 0.07$; Left FFA: $p = 0.26$; Left PPA: $p = 0.54$; Right PPA: $p = 0.44$). We then tested whether the category-level reactivations in right HPC and right FFA was significantly associated with subsequent (1-week-delayed) recognition (total correct). We found no significant relationship between recognition accuracy and category-level reactivations in right HPC ($F(32,60) = -0.27$, $\beta = -0.15$, $SE = 0.55$, $p= 0.79$). On the other hand, we found an inverse relationship between category-level reactivations and recognition in right FFA ($F(32,60) = -3.39$, $\beta = -2.37$, $SE = 0.7$, $p= 0.001$). Importantly, this effect was not significant in pre-encoding rest period ($p = 0.29$), suggesting that the inverse relationship is experience dependent. Together, these results replicate our main findings with the event-specific reactivations.

We next tested HPC-cortex co-reactivations using the category-level patterns described above. We did not any significant between post- versus pre-encoding rest differences for HPC-FFA ($p = 0.14$) or HPC – PPA ($p = 0.9$) co-reactivations. Thus, we did not test any memory effects for co-reactivations of category-level patterns.
